# Analysis of Naturally Occurring Somatic Insertions in the Human Genome

**DOI:** 10.1101/2025.09.24.677890

**Authors:** Chih-Lin Hsieh, Zarko Manojlovic, Timothy Okitsu, Cindy Okitsu, Jordan Wlodarczyk, Nick Shillingford, Ramzi Bawab, Yong Hwee Eddie Loh, Michael R. Lieber

## Abstract

Biochemical and genetic experimental systems permit precise definition of enzyme requirements and mechanistic steps in DNA repair. Comparison of these findings to repair events at naturally occurring breakage sites in multicellular organisms is valuable for confirming and extending these insights. However, heterogeneity in any cell population increases with each cell division, and the reliable detection of DNA breakage sites and their repair *in vivo* has been difficult due to technical limitations. Here, we examine somatic insertional mutations naturally occurring during normal metabolism and cell division in single human colon crypts using a novel whole-genome sequencing method. We find that replication slippage is a dominant mechanism for these events, and insertions larger than 10 bp are uncommon. Mechanistic features of these sites in physiologically normal cell clones, such as single human colon crypts, permits inferences about the DNA breakage repair zone and processing within natural chromatin, thereby permitting comparisons to experimental studies using *ex vivo* cellular and simplified biochemical systems.

## INTRODUCTION

Insertional mutations in human somatic cells can derive from a variety of causes. Strand slippage during DNA synthesis, either replicative synthesis during S-phase or at a DNA breakage site, can give rise to insertions. In these strand slippage events, adjacent sequence is duplicated leading to 1) extra full or partial unit(s) of the repeat being inserted if the slippage occurs within a previously existing repetitive sequence or 2) a new direct repeat if the slippage occurs within a unique sequence. When novel nucleotides without any relation to the surrounding sequences on the same strand are inserted, DNA breakage must occur prior to the insertion.

Both single-strand breaks (SSBs) and double-strand breaks (DSBs) can arise from many hydrolytic, oxidative, and enzymatic events during physiological DNA and cellular metabolism (1). Base excision and nucleotide excision repair handle most SSB sites. There are four repair pathways for DSB repair in eukaryotic cells (2, 3). The two main pathways are homologous recombination (HR), which occurs predominantly during S-phase, and nonhomologous DNA end joining (NHEJ), which takes place in all phases of the cell cycle. Single-strand annealing (SSA) and alternative end joining (aEJ) are two additional pathways that function primarily during DNA synthesis.

DNA breakage repair analysis in experimental systems has several complications that limit the extrapolation to naturally occurring events in human somatic cells. Cell-free biochemical studies use naked DNA substrates that typically lacking histones, which can have a large effect. The ratio of relevant DNA repair enzymes and factors in biochemical reconstitution experiments is also not the same as in human cells in culture or within the human body. In many experimental systems, exogenous substrates are introduced in large numbers simultaneously that can overwhelm repair systems and introduce potential complications from other molecules. This is also true for cell-free biochemical systems as well as substrates introduced into the cells (often into the cytoplasm rather than the nucleus) in transfection studies. Experimental systems often focus on very limited zones or artificially introduced DNA substrates and limit extrapolation to physiological conditions and naturally arising events. In all types of studies, if the DNA breaks are created at slightly different points in time or distant locations in the genome, then the two DNA ends may be far apart and initially processed independently prior to being brought into proximity for any final phase of repair.

Despite the enormous amount of human genomic sequence information generated to date, actual quantitative and qualitative analyses of short insertion events (<50 bp) in somatic cells using WGS has been very limited for several reasons. First, most somatic cell sampling methods contain a population of cells that diverged many divisions before the sample was taken. New distinct mutations arise in each cell division and only on one of the two alleles. These distinct mutations are only present at very low allele frequency in the cell population examined and cannot be reliably detected by WGS due to the low count of alternate (mutant) allele sequencing reads. The consensus sequence from WGS of a somatic cell sampling is the genome of the common progenitor shared by all the cells in the cell population. Therefore, there is little capability in identifying somatic mutations beyond the diploid fertilized egg genome that gave rise to that individual.

Second, efforts to delimit the complexity of human WGS samples have complications. Single cell WGS is an approach to identify heterogeneity. However, the need for whole genome amplification (WGA) and the incomplete efficiency of adaptor ligation in the WGS library construction can introduce DNA mutations as well as coverage bias (4–7). *Ex vivo* culture and expansion of single cells is another approach. However, a clonal population of cells derived from *ex vivo* culture of a single cell can acquire mutations induced by culturing conditions.

The epithelial lining of the human colon consists of over 10 million colon crypts. The ∼2000 cells that line each human colon crypt derive from one or very few stem cell lineages. Therefore, these ∼2,000 cells are natural cell clones and can be used to identify somatic DNA changes easily and reliably with a sequencing library constructed directly from a single colon crypt without DNA extraction and without WGA (8).

DNA breakage repair often leads to DNA sequence change beyond single base substitution. Here, we identify DNA breakage repair events among simple insertions to compare with findings from experimental systems. Naturally occurring insertions derived from DNA breakage repair events are not likely to be duplicated in different individuals. In the process of identifying DNA breakage repair events, we discovered that hidden inter-sharing of somatic insertion variants in unrelated individuals is very common. Importantly, this can lead to a false interpretation of a variant as a true positive somatic variant. These false positive variants may overwhelm the true positive variants in some applications of WGS analysis. Our experience and solution for identifying the false positive variants will be useful for others in the future (see Methods).

The ability to reliably identify somatic mutations in natural cell clones permits examination of physiological DNA breakage and repair events in the human body. Here, we analyze the simple insertion variants identified by conventional somatic variant callers in high depth WGS of 106 single human colon crypts from 21 individuals to explore the occurrence and appearance of DNA breakage events. We are particularly interested in deepening our understanding of DNA breakage events and the DNA repair process in their natural physiological context, which has native chromatinized DNA and DNA repair enzymes in their normal nuclear ratios.

## MATERIALS AND METHODS

### Tissue collection, crypt isolation, sequencing library construction, and whole genome sequencing

A small piece of normal colon is collected through the Pathology Department Tissue Procurement Core with an approved IRB from individuals who have undergone surgery to remove part of the colon under the standard of care at either Keck Hospital of USC or Children’s Hospital Los Angeles. Tissue processing, crypt isolation, and WGS library construction have been described in detail previously (8).

A total of 106 crypt libraries and 21 bulk libraries are constructed from 21 individuals of age 10 months to 90 years old. After quality control, libraries are pooled and sequenced on seven S4 flow cells using NovaSeq 6000 (Illumina, San Diego) S4 300 cycles reagent kit (v1.5) in the Keck Genomics Platform (KGP) Core facility at USC. The post-alignment statistics of 91% > 20x and quality measures of these libraries are provided in a previous publication (8).

### Variant calling

The sequencing reads are assessed and analyzed using a validated in-house pipeline (based on GATK v4.2 best practices) from the KGP as described previously (8) and the DRAGEN Somatic v 4.2.4 pipeline (Illumina Inc) after sequencing runs. PCR duplicate reads were identified and marked using Picard (MarkDuplicates) as part of our standard GATK preprocessing pipeline. These marked duplicates were excluded from downstream variant calling steps to prevent artificial inflation of variant support. Two germline callers, HaplotypeCaller and Freebayes (v1.2.0), and two somatic variant callers, Mutect2, Strelka (v2.9.0), are used in the KGP pipeline. Somatic variant calls in the crypt samples are made with the matched bulk tissue sample from the same individual as the control. The shared somatic variant calls from Mutect2 and Strelka (2Caller) through the KGP pipeline and DRAGEN pipeline have a high inter-caller concordance with 93.78% (s.d. 2.79%) 2Caller calls present in DRAGEN and 80.77% (s.d. 4.92%) DRAGEN calls present in 2Caller. The variant calls from 2Caller are used to increase variant call confidence for all downstream analyses. The variant calls from Illumina’s DRAGEN somatic pipeline are only used for internal validation of the analysis and are not used in the downstream analysis.

### Analysis of Simple insertions

Insertion can occur in three ways: 1) strand slippage during DNA replication due to misalignment of repetitive sequences during DNA replication, resulting in repeat expansion; 2) strand slippage or DNA breakage repair in an unique DNA region, creating a new direct repeat; 3) DNA breakage repair leading to insertion of novel nucleotides without substantial consecutive nucleotides matching to the surrounding sequences on the same DNA strand. One of the goals here is to identify DNA alterations derived from the repair of DNA breakage instead of DNA replication errors that occur without DNA breakage. Therefore, we are particularly interested in insertions involving novel nucleotides added at the site of DNA alteration.

A total of 16,483 autosomal insertion variants are identified in 106 crypts (21 individuals) by the 2Caller (Fig. 1). First, duplicate variant calls present in more than a single individual (apparent inter-sharing) are removed from the 16,483 insertions because identical somatic DNA breakage events are unlikely to occur across different individuals. Second, only one entry is kept for duplicate variant calls from different crypts of a single individual (intra-sharing).

**Figure 1.**
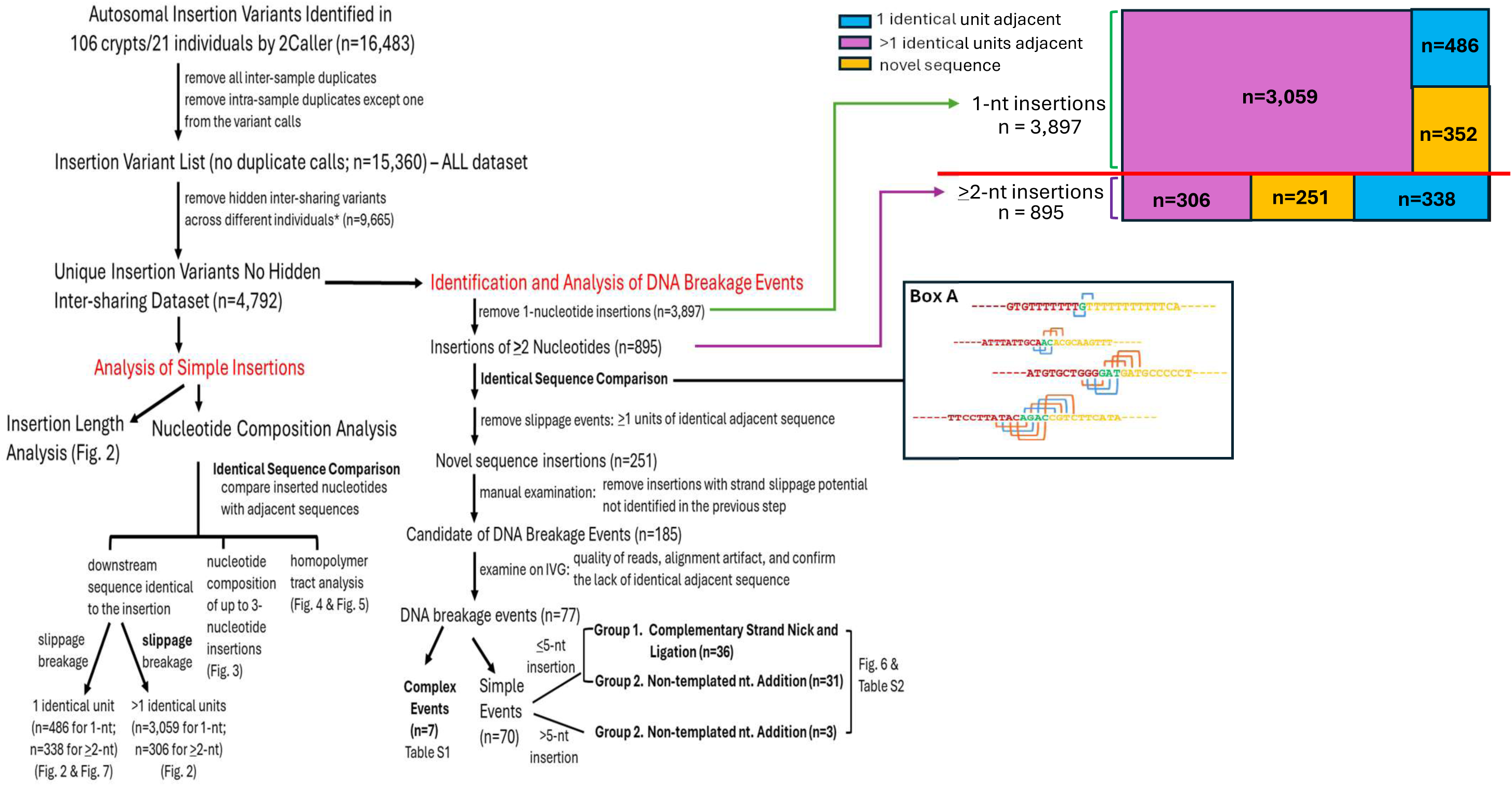
Analysis Flow Chart for Simple Insertions. Insertion events likely due to strand slippage and DNA breakage repair are separated. **Box A**, examples from the observed insertions illustrating the identical sequence comparisons are presented with red letters for upstream sequence, green letters for inserted sequence, and yellow letters for downstream sequence. Orange bracket indicates base-to-base match and blue bracket indicates the lack of base-to-base match.

The variants with no apparent duplicates are further analyzed (n=15,360, ALL dataset). In our analysis of the ALL dataset, we discovered “hidden inter-sharing” events that are not apparent in the 2Caller variant calls. These hidden inter-sharing events are only called in one or more crypts from a single individual, however, they actually are present in more than a single individual. These hidden variants can be present in both the crypt and bulk control (therefore, not called as a somatic variant) in other individuals. Also, the sequencing reads for the alternate allele do not meet the variant calling threshold (therefore, not a “PASS” variant) in other individuals. These hidden inter-sharing variants appear to only occur in a single individual and are falsely concluded as unique variant calls. To strategically filter for these hidden inter-sharing variants, we utilize VCFs from Freebayes (v1.2.0). Freebayes is a haplotype-based caller commonly used for germline variant detection in a multi-sample dataset. Freebayes reports the variant location in all samples with alternate allele sequencing read(s) in the dataset when the variant call of a putative alternative allele is supported by the sequencing reads in any sample. We developed a script to identify these hidden inter-sharing events in the Freebayes output VCF files (https://github.com/twewyttst777/VariantDuplicateCheck) to further remove variants with hidden inter-sharing from the 2Caller insertion variant list of 15,360. However, matching variants from a somatic caller (Mutect2 or Strelka) to FreeBayes can be challenging due to differences in variant representation, calling logic, and filtering thresholds. Additionally, indels and complex variants may be left-aligned differently or split/merged in inconsistent ways across tools, making direct position-based comparison error-prone. Of the 15,360 variants, 904 cannot be matched in Freebayes VCFs. We chose to exclude these 904 variants from the final analysis to ensure that no inter-sharing variants across individuals are in the analysis. After removing the variants with hidden inter-sharing, a total of 4,792 unique insertion variants is further analyzed.

The sequences of 50 nucleotides immediately upstream and downstream of each of the 4,792 unique insertion variants were extracted from the GRCh38 reference genome for further analysis. Each base of the inserted nucleotides is compared base-to-base to the adjacent sequences for an exact match of the entire insertion (identical sequence comparison, Fig. 1). As expected, no exact match to the insertion was found immediately upstream of the inserted sequences, consistent with the behavior of alignment algorithms that left-align indels according to VCF conventions (9). Therefore, only the sequence immediately downstream of the insertion is further analyzed. The length of each insertion is counted, and insertions with one or more units of identical sequence immediately downstream of the insertion (not counting the insertion as a unit) are identified and counted. Simple sequence repeats (SSR) of 2 to 6 nucleotides downstream of the insertion are identified by identical sequence comparison followed by manual inspection and counted for ≥2 nucleotide insertions. Homopolymer length is counted for 1- to 3-nucleotide insertions in a homopolymer tract immediately downstream of the insertions.

### Identification and analysis of DNA breakage events

To identify naturally occurring DNA breakage events in the human somatic genome, a very stringent set of criteria is applied to filter out variants that can potentially derive from replication slippage. In the three ways an insertion can occur described above, only #3 is without any strand slippage. First, all 1-nt insertions are removed from the 4,792 unique insertion variants because it is difficult to conclusively determine the mechanism for 1-nt insertion. In the second step, variants with one or more units of identical sequence to the inserted nucleotides immediately downstream from the insertion site are excluded from the 895 ≥2-nt insertion variants for breakage event identification. The remaining 251 novel sequence insertion events are further examined manually to remove insertions that are the product of a nearby repeat expansion that the identical sequence comparison cannot identify. The final 185 insertions that are potential candidates for DNA breakage events are further examined on IGV to remove variants that have any potential to be the result of slippage. When the inserted nucleotides have no substantial sequence relations beyond one or two consecutive nucleotides to the adjacent sequence on the same DNA strand, breakage of the DNA strand must occur to allow insertion of nucleotides. We find a total of 77 insertion variants that are clearly the product of repair at DNA breakage sites without evidence of sequence relationship to the surrounding sequences. Although PASS variants from Mutect2 and Strelka clearly meet the threshold for variant calling with multiple support alternate (mutant) allele reads and good quality scores, the minimum mapping quality is examined to confirm the quality of the calls in addition to manual examination on IGV. These 77 insertion variants are further categorized into three subgroups: complex events; partially complementary insertion (Group 1); and events with entirely non-templated nucleotide additions (Group 2).

## RESULTS

### Analysis Strategy

A total of 16,483 somatic insertion variants on the autosomes in 106 single colon crypts from 21 individuals are called by both Mutect2 and Strelka (2Caller) with a matched bulk tissue control for each individual. Only the common PASS calls from Mutect2 and Strelka are included in the insertion variant list to ensure the variants identified are of high quality for our analysis of naturally occurring insertions. Variants called repetitively in different individuals (inter-sharing) are more likely to be germline variants (despite a matched bulk tissue control), mutation hotspots, or alignment artifacts. Also, variants shared by multiple crypts from the same individual (intra-sharing) most likely only occurred once in the individual prior to the separation of these crypts that share that specific somatic mutation during tissue development. Therefore, inter-sharing variants and duplicate entries of the intra-sharing variants are removed to generate a dataset of 15,360 variants for further analysis (ALL dataset, Fig. 1). In our analysis of these 15,360 variants with no duplicate variant calls, we discovered that hidden inter-sharing events are quite common among these variants that appear to be without duplicates and present only in a single individual. The hidden inter-sharing events have alternate allele sequencing reads present in multiple individuals but are only called as a somatic variant in a single individual by Mutect2 and Strelka. These hidden inter-sharing events are predominantly false positive calls due to alignment issues, borderline local read counts, and lower mapping quality. A final count of 4,792 (30.7% of the 15,360 total insertion variants from 2Caller) unique insertion variants, after removing the hidden inter-sharing variants using a script that we developed (see Methods), are used for further analysis (called the “no hidden inter-sharing dataset”).

### Large naturally occurring somatic insertions are uncommon

Among the total of 4,792 unique insertion events in the no hidden inter-sharing dataset, single base insertions are the most abundant (n=3,897; ∼81% of the total) and the event count decreases with increasing insertion length (Fig. 2A, 2B, and 2C). Distribution of the insertion length is very similar for the ALL dataset of 15,360 variants and the no hidden inter-sharing dataset of 4,792 variants (compare Fig. 2 and Supplementary Fig. S1). Detection of insertion length is limited to <50 bases by the 2caller because Strelka has a default indel size limit of 49. The majority of the insertions are less than 10 nucleotides in length (Fig. 2A, 4725/4792 = 98.6%) as previously reported by others (10). The longest insertion length observed is 43 in the no hidden inter-sharing dataset (Fig. 2A). This finding indicates that large somatic insertions are uncommon in the human somatic genome.

**Figure 2.**
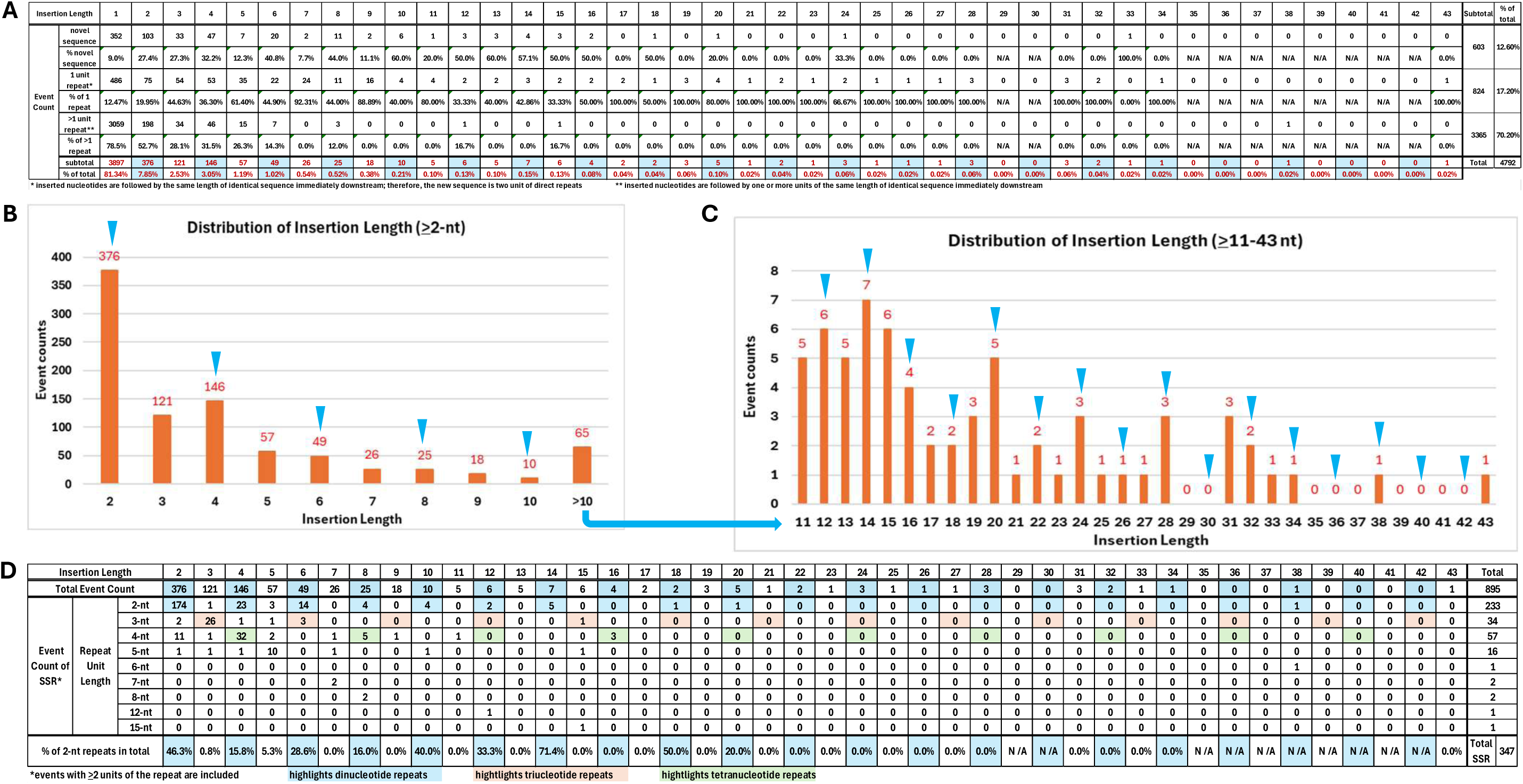
Distribution of Insertion Length. A) Event count and the observed frequency in the total for each insertion length in the no inter-sharing dataset of 5,695 insertions from 106 single human colon crypts (21 individuals). The count of insertions with no identical duplicate sequence (novel sequence), a single unit of duplicated sequence, and ≥1 unit of duplicated sequence in the surrounding region for each insertion length is also listed. Percentage of events with ≥1 unit of duplicated sequence in total occurrence of each insertion length is also calculated. The blue highlight marks the even number insertion events. B) Graphical illustration of the event counts for insertion length from 2 to 10. C) Graphical illustration of the event counts for insertion length from 10 to 45. For B) and C), the counts are displayed on the top of each bar with blue arrows marking the even number nucleotide insertions. D) Simple sequence repeat (SSR, length of 2 to 15) with at least 2 units of repeat adjacent to each insertion length is identified and listed. The percentage of 2-nt SSR in the total for insertion length is calculated. The highlights are as indicated.

### Naturally occurring somatic insertions are predominately replication slippage events

The identical sequence comparison reveals that ∼87% (824 + 3365 in a total of 4,792) of the insertions have at least one unit of duplicate (identical) sequence immediately downstream of the insertion (Fig. 2A; see ALL dataset in Supplementary Fig. S1). It is important to note that the left-align preference of the software always positions the duplicate sequence immediately downstream, even though the actual insertion could be any unit within a series of repeats. When a single unit of duplicate sequence is downstream from the inserted nucleotides, the insertion creates a direct repeat at the previous unique sequence site. When there is more than a single unit of duplicate sequence downstream from the inserted nucleotides, the insertion expands the unit of previously repetitive sequence.

As mentioned above, the insertion event count decreases with insertion length in general. Interestingly, insertion events with an even number of nucleotides inserted appear to occur more frequently than events with one less nucleotide inserted (Fig. 2A, compare insertion length marked by blue highlight to the event count without highlight immediately to its left; Fig. 2B and 2C, see blue arrowheads). We hypothesize that this phenomenon is due to the abundance of the dinucleotide repeats in the human genome (11–15). An analysis of SSR presence downstream of the insertions shows that the SSR, especially dinucleotide repeats, accounts for a high fraction of the insertion events (Fig. 2D). This analysis demonstrates that the abundance of the SSR in the human genome correlates with the presence of insertion events in human somatic cells, most likely due to replication slippage being the primary mechanism of insertion.

### Inserted nucleotides do not always reflect the overall genomic nucleotide composition

The nucleotide composition in the human genome is about 60% A/T nucleotides and 40% C/G nucleotides (12). Single nucleotide (1-nt) insertion is the most frequent insertion event (3,897 total) in our study (Fig. 3A). Among 1-nt insertions, A/T nucleotides account for 87.2% (42.0% A + 45.2% T) while C/G nucleotides account for 12.8% (6.2% C + 6.6% G) of the total events (Fig. 3A; Supplementary Fig. S2A for ALL dataset). Among the 376 two-nucleotide (2-nt) insertions, the nucleotide composition for the first inserted base is 80% A/T (39.1% A + 41.0% T) and 20% C/G (10.4% C + 9.6% G). The nucleotide composition for the second inserted base is 72.6% A/T (35.4% A + 37.2% T) and 27.4% C/G (13.6% C + 13.8% G) (Fig. 3B; Supplementary Fig. S2B for ALL dataset). A total of 121 three-nucleotide (3-nt) insertions are identified, with AAA, CAG, AAT, AGA, TTA, and TTT being the most abundant (Fig. 3C; Supplementary Fig. S2C for ALL dataset). While A/T nucleotides are mostly over-represented in the 1- to 3-nt insertions compared with the A/T content of the human genome, the representation of the third base for the 3-nt insertions is much less biased (Fig. 3; Supplementary Fig. S2 for ALL dataset).

**Figure 3.**
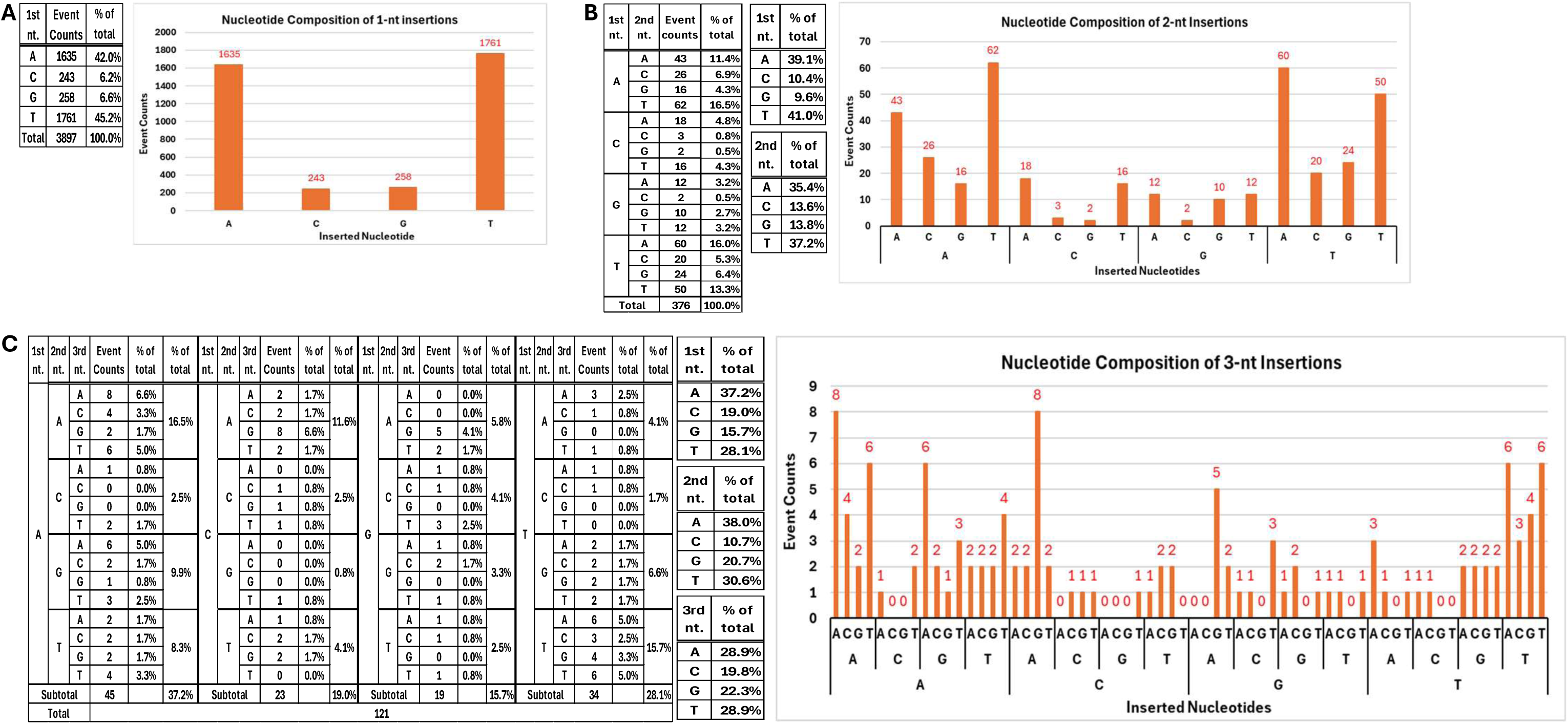
Nucleotide composition analysis of one to three nucleotide insertions. A) Nucleotide composition of single base (1-nt) insertions. B) Nucleotide composition of two-base (2-nt) insertions. C) Nucleotide composition of three-base (3-nt) insertions. The tabulation of nucleotides and percentage of each of the four nucleotides at each position of insertion is on the left and the graphical illustration of the nucleotide profile with event count on the top of each bar is on the right of each panel.

### The occurrence of insertions within the homopolymer tracts is not correlated with the presence of homopolymer tract length in the human genome

Among the 3,897 1-nt insertions, 3,545 are followed by a stretch of 1 to 13 nucleotides identical to the inserted nucleotide (homopolymer) with 1, 6, 7, and 8 being the most abundant homopolymer lengths (Fig. 4A; Supplementary Fig. S3A for ALL dataset). Of the 106 2-nt insertions with two identical nucleotides inserted, 93 are followed by homopolymer lengths of one to seven (Fig. 4B; Supplementary Fig. S3B for ALL dataset). Of the 16, three identical nucleotide insertions, 12 are followed by homopolymer tracts of one to six in length (Fig. 4C; Supplementary Fig.S3C for ALL dataset).

**Figure 4.**
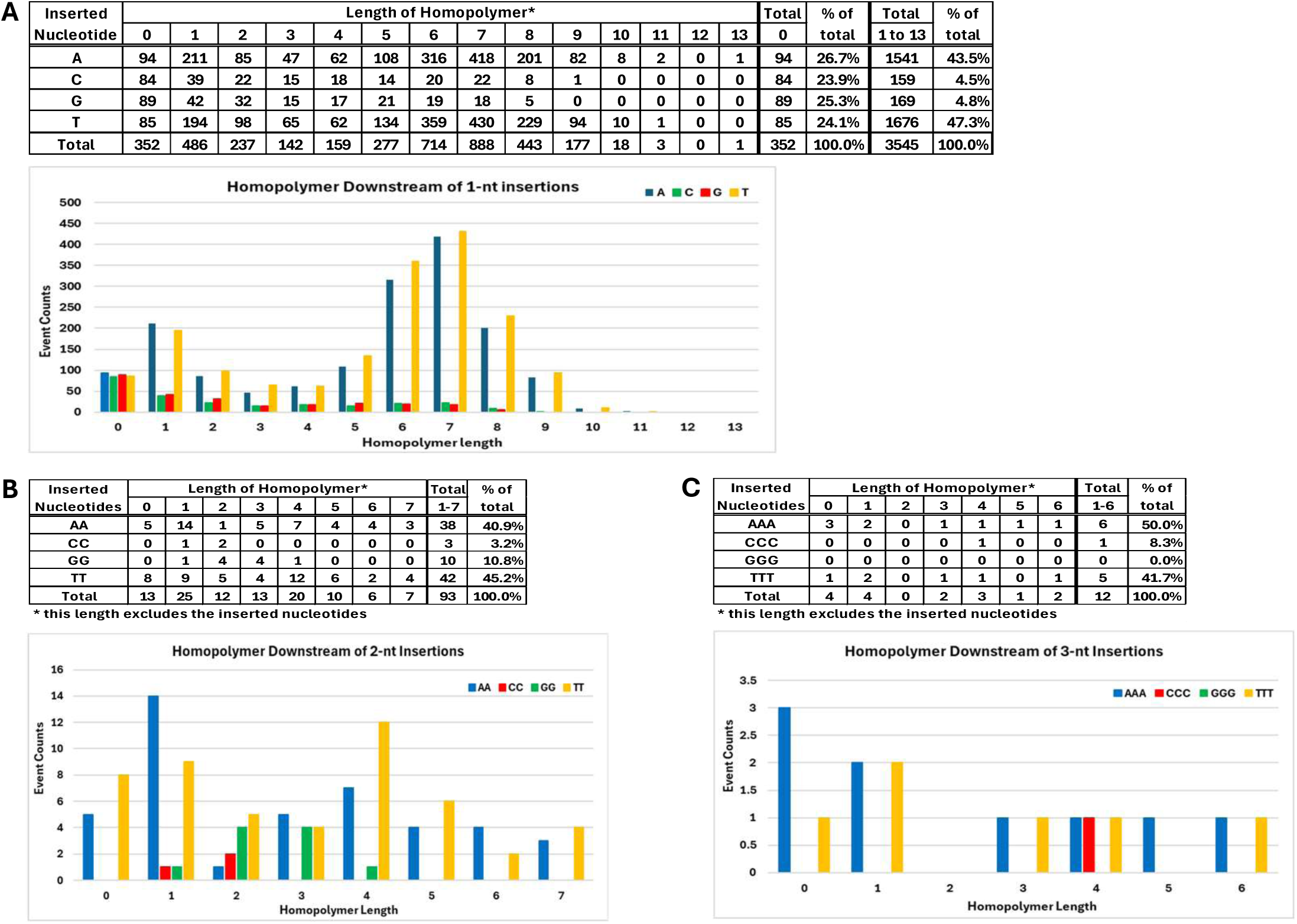
Presence of homopolymer at site of one- to three-nucleotide insertions. A) Count of events with homopolymer length 0 to 12 at 1-nt insertion sites. Length of 0 indicates the 1-nt insertion is followed by a different nucleotide than the base inserted. B) Count of events with homopolymer length 0 to 9 at the site of 2-nt insertions with two identical nucleotides. C) Count of homopolymer length 0 to 6 at the site of 3-nt insertions with three identical nucleotides. The tabulation of events is above the graphic illustration in each panel. All lengths of homopolymer present in the dataset is included in the illustration.

In the human genome, the abundance of homopolymer tract decreases with the homopolymer tract length, and A/T tracts are ∼1.4-fold more abundant than C/G tracts (Fig. 5A) (16). Interestingly, the occurrence of insertion in A and T homopolymers of 6 and 7 in length are more frequent than in the length of one as mentioned above. When the relative abundance of the homopolymer tracts observed in the 1-nt insertions is compared with the reported frequency of homopolymer length in the human genome, the A/T homopolymer lengths of 6 to 9 are clearly overrepresented in the insertional events (Fig. 5B). Likewise, a dramatic over representation of the 6 to 9 nucleotides long G/C homopolymers is observed (Fig. 5B). This analysis demonstrates that the insertions, at least the 1-nt insertions, occurring in the homopolymers do not reflect the relative abundance of different lengths of homopolymer in the human genome. It is curious how the mechanism of slippage within homopolymer tracts during replication leads to over representation of homopolymer lengths of 6 to 9.

**Figure 5.**
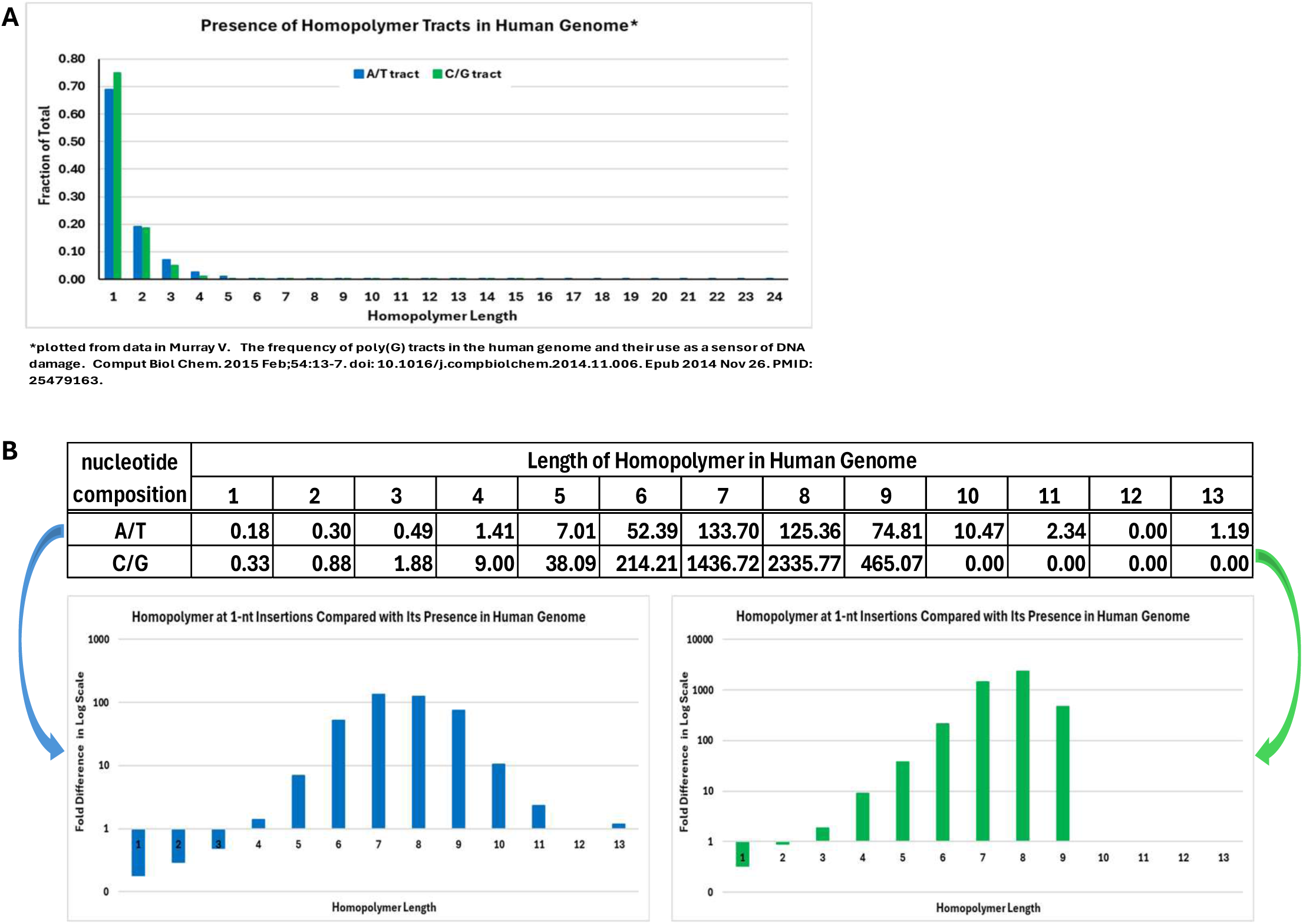
Presence of homopolymer at 1-nt insertion sites. A) Abundance of different lengths of homopolymer in the human genome. B) Fold difference of the observed frequency in our dataset and the frequency of the same homopolymer length in the human genome, plotted with logarithmic scale (see Table above for actual fold difference). Group 1. Complementary Strand Nick and Ligation (n=36)

### DNA breakage repair events

As a reminder, 1-nt insertions and all apparent and hidden inter-sharing variants have been removed prior to this analysis. There are 895 events with ≥2-nt inserted, and 251 of these events do not have identical sequence immediately downstream from the insertion (Fig. 1). Upon manual examination of 50 nucleotides upstream and/or downstream of each insertion, insertions that are clearly due to SSR expansion but not identified in the identical sequence analysis are removed from the list of 251 insertions. The remaining 185 novel sequence insertions are then further examined on IGV for any clear signature of DNA breakage to account for the event. A total of 77 insertion variants have no discernible evidence of sequence relation to the surrounding sequences on the same strand on which novel nucleotides can be inserted by any conceivable mechanism without DNA strand breakage (Fig. 1). We used the MMQ field (median mapping quality) to assess the read alignment quality. In these 77 insertions, all the alternate allele-supporting sequencing reads have an MMQ ≥30 (with 73 having the score of ≥60 (Supplementary Table S1 and S2), indicating high-confidence alignments.

Among these 77 DNA breakage events, seven involve a more complicated breakage/repair process beyond simple insertion. In these events, the minimal footprint of the DNA damage (minimum repair zone) can be inferred from the outermost DNA changes in the vicinity. These complex events, with a combination of deletion, insertion, and single nucleotide substitution, are likely the result of NHEJ repair (Supplementary Table S1). It is noteworthy that two of the variants, a 2-nt insertion and a 4-nt insertion with three nucleotides in between, are from a much more complex repair event at a common breakage site (Supplementary Table S1 and S2, marked with red # at the end of their Chr_POS).

Among the remaining 70 simple insertions of novel sequence, 67 have ≤5 nucleotides inserted. We categorize these 67 short insertions into two groups: Group 1, a substantial stretch of consecutive identical nucleotides to the inserted nucleotides are found on the complementary strand at the insertion site; Group 2, no sequence relationship between the inserted nucleotides and adjacent sequences (Supplementary Table S2). It is always possible that nucleotides are randomly inserted without any relations to the adjacent sequences at a DNA breakage site no matter how unlikely it may be. Alternatively, the 36 variants in Group 1 can be the result of a nick and ligation process such that a short DNA fragment is released due to an endonuclease nick near a double strand DNA break, and then followed by ligation of the fragment to the end of the complementary strand at the double-strand break (Fig. 6A; Supplementary Table S2). When this nick and ligation process occurs after a DNA polymerase adds one or more nucleotides to the broken end, the added nucleotides would be present at the junction (Fig. 6B; Supplementary Table S2). There are 31 variants with inserted DNA fragments with no sequence relation to the adjacent sequence and are categorized as Group 2 (Fig. 6C; Supplementary Table S2). Three final variants have a much longer insertion with one harboring a five consecutive nucleotide match to the downstream sequence three bases away from the DNA break junction, while the other two have a completely novel sequence inserted (Supplementary Table S2).

**Figure 6.**
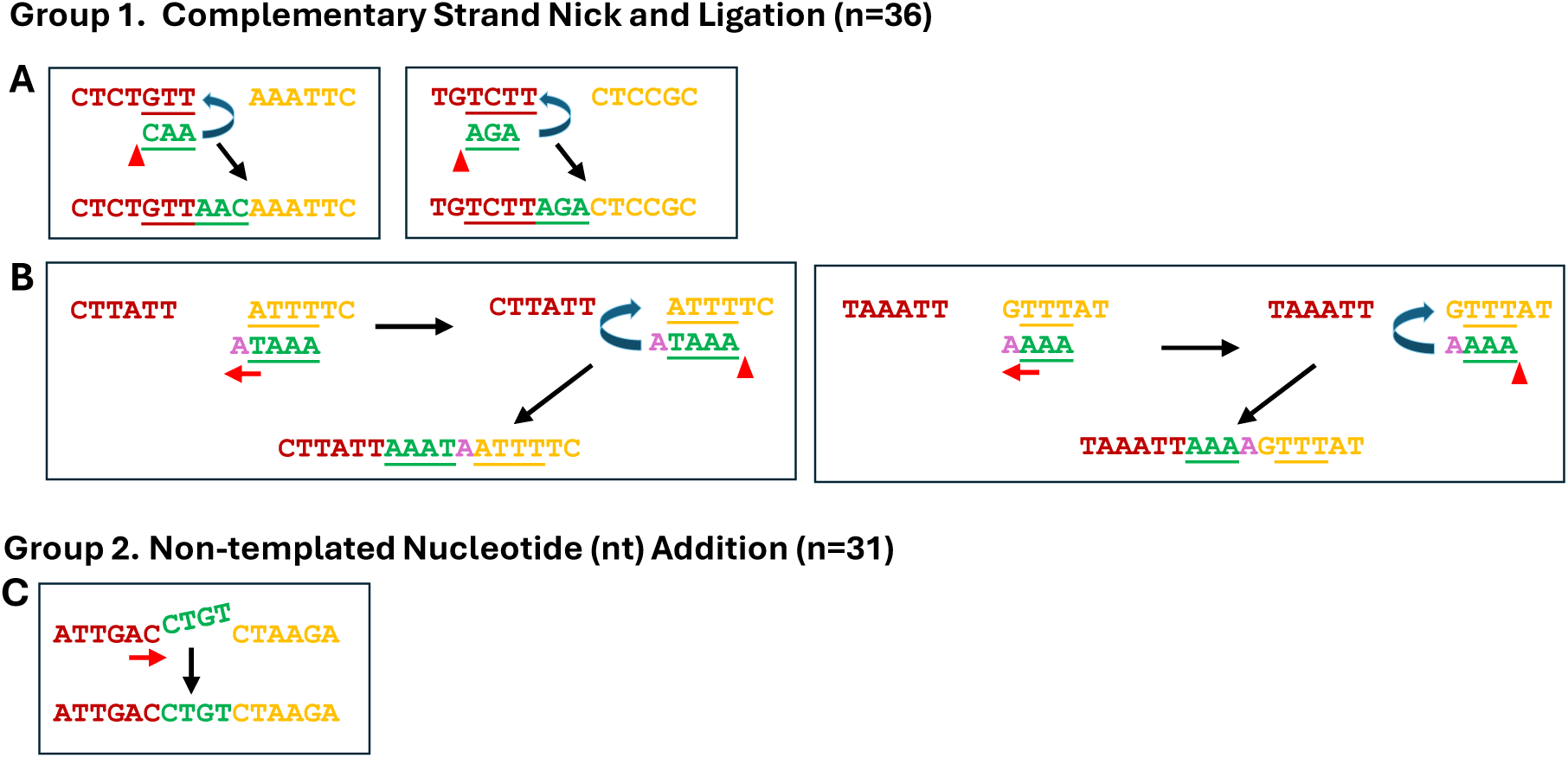
Two Groups of Simple Insertions ≤5 in length illustrated with observed naturally occurring insertions. In addition to random nucleotide addition as the mechanism, panels A and B illustrate possible non-random mechanisms. Group 1, nick and ligation, consists of inserts with at least partial homology to the complementary DNA strand illustrated in examples from our dataset A) with no additional nucleotide inserted and in B) with extra nucleotide inserted. Group 2, Non-templated random nucleotide addition: C) no apparent template in vicinity for inserted nucleotides with illustration of an event observed in our dataset. See Supplementary Tables S1 and S2 for information on all events from our dataset. The red letters Indicate nucleotides immediately upstream of the insertion, the yellow letters depict nucleotides immediately downstream from the insertion, green letters mark the inserted nucleotides, and the pink letter marks addition of a new nucleotide. The underlined letters are the complementary sequences.

### Insertion with one unit of duplicated sequence in non-repetitive DNA region

Insertion of a single unit of duplicated sequence in a unique DNA region will result in a new direct repeat, while a single unit of duplicated sequence in a repetitive region will be a repeat expansion. In a repetitive DNA region, microhomology between the 3’-end of the insertion and the nucleotides upstream of the template strand can facilitate mispairing. In contrast, strand slippage in a unique DNA region does not always have microhomology present to mediate the mispairing. In the total of 338 insertions ≥2-nt that generate a new direct repeat, 150 have no microhomology that could mediate the mispairing (Fig. 7A). Among the remaining 188 events, 179 have 1 to 7 bases of microhomology to mediate the mispairing, and 9 are of a complex nature or have random base changes within the duplicate (Fig. 7A).

**Figure 7.**
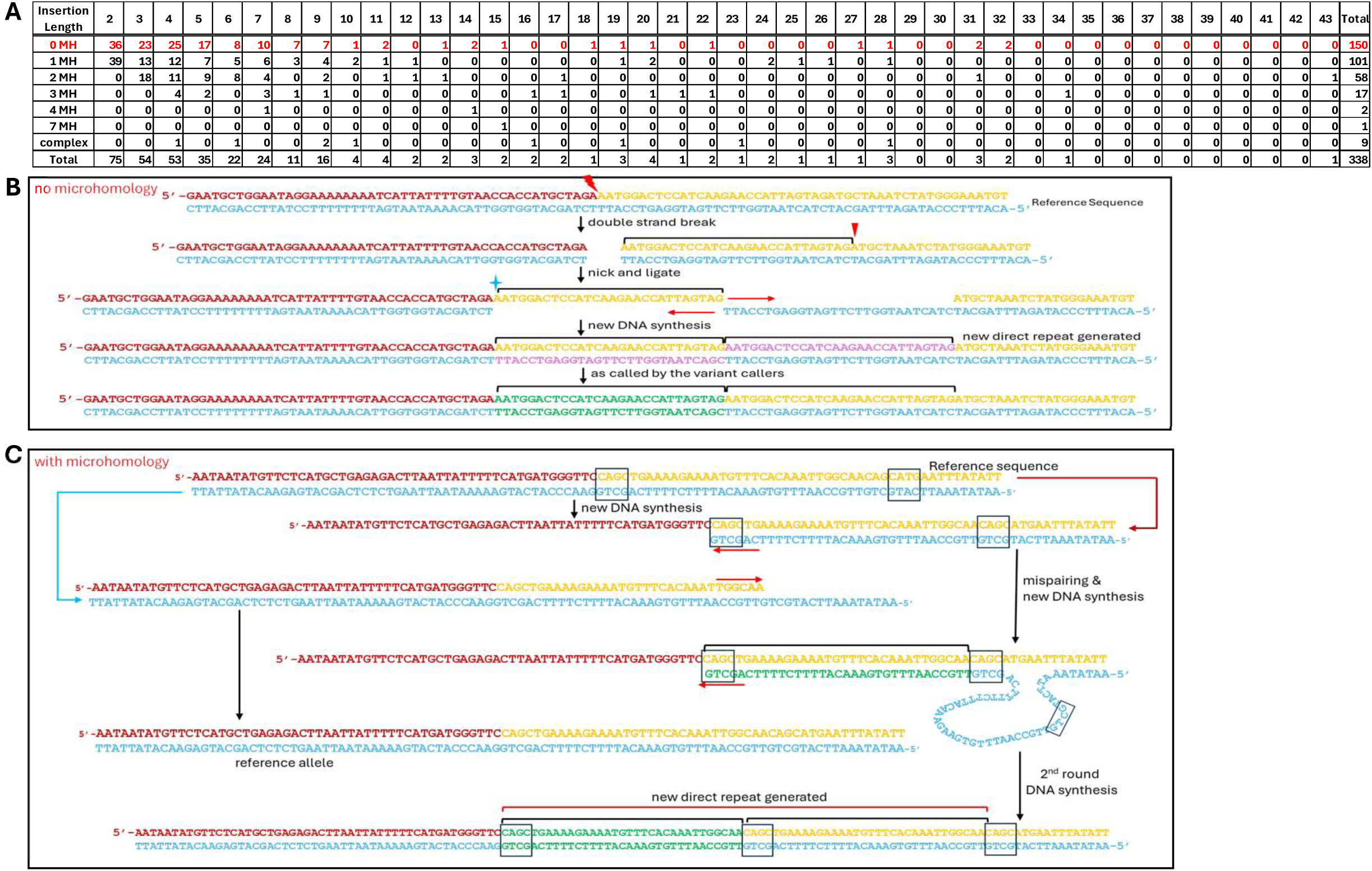
Potential Pathways to Generate Insertions of a Single Unit Duplicated Sequence. A) Presence of microhomology at the insertion site of the 372 insertion events of ≥2-nt with a single unit duplication in our dataset. MH = microhomology. For the same strand nick and ligation pathway as illustrated in B), the red jagged flash mark indicates the site of a double strand break, the red vertical arrowhead indicates the site of a single strand nick, the blue star indicates the site of ligation, and the red arrows indicate the direction of DNA polymerization. For the DNA slippage with microhomology pathway as illustrated in C), the markings are as described above, and the microhomology involved in the sequence duplication is marked by a rectangle box.

## DISCUSSION

Our study finds that 1) insertions of >10-nucleotide are uncommon in human colon crypts; 2) replication slippage is the primary mechanism for naturally occurring somatic insertions; 3) DNA breakage repair is a likely cause of only a very small number of somatic insertions; 4) simple sequence repeats (SSR), especially dinucleotide repeats, represent the most abundant events in the even-numbered nucleotide insertions; 5) representation of the four nucleotides, A, C, G, and T, in the insertions is quite different from that in the human genome; and 6) homopolymer tracts account for a high fraction of the somatic insertions and homopolymer lengths of 6 to 9 bp are over-represented among the insertions in the homopolymer tracts in the human genome. We categorized the insertions caused by DNA breakage repair into two major groups with proposed potential repair pathways (Fig. 6). We also proposed potential pathways for insertions that generate a new direct repeat with a single unit duplication, which accounts for ∼37.8% of the >2nt insertions (338 events among 895, and Fig. 7B and Fig. 7C).

It is not likely that any of our observations are due to sequencing errors or the 10 cycles of PCR in the library enhancement step. Random PCR-induced errors are unlikely to recur at the same genomic positions across multiple molecules and independent libraries. Therefore, random PCR-induced errors are not expected to be consistently identified as somatic variants by standard variant callers. Non-random PCR artifacts due to the nature of DNA sequences can occur repeatedly in multiple crypt samples and in bulk DNA controls of different individuals. A DNA change must occur repeatedly on a substantial number of molecules in the reactions for some colon crypt samples and not in the bulk DNA for it to pass the threshold to be called by the somatic variant callers. All the variants in our 2caller (common calls from Mutect2 and Strelka) variant list are supported by multiple good quality unique sequencing reads (after duplicate reads removal), which passed the calling threshold and are only present in the colon crypts and not in the bulk DNA control of each individual. In addition, we remove all inter-sharing variants (both those that appear on the variant list and those that are hidden) in more than a single individual among the 21 individuals. Therefore, most, if not all, of the potential artifacts are eliminated.

Overall, an average of 45 somatic insertions is identified per colon crypt (4,792/106 = 45.2) and the event count decreases with insertion length in general. Insertions >10 nucleotides only account for 1.3% of the somatic insertions. It is noteworthy that SSR only accounts for 3% of the human genome (14), but 38.8% (347/895, Fig. 2A) of the insertion events take place within SSR regions. The presence of insertions primarily reflects the abundance of dinucleotide SSR in the human genome. A majority of the naturally occurring insertions are in either homopolymer tracts or SSR, most likely due to strand slippage during DNA synthesis, even though the possibility of DNA strand breakage cannot be entirely ruled out. We infer that sites of insertion due to DNA breakage do not occur very frequently.

While the presence of homopolymer tracts decreases with homopolymer length in the human genome, the occurrence of 1-nt insertions at homopolymer tracts lengths 6 to 9 are vastly over-represented. At the same time, the lengths of 1 to 3 are slightly under-represented compared with their presence in the human genome (Fig. 5B). The number of 2-nt and 3-nt insertions occurring in homopolymer tracts is too small to make any inference about tract length representation. It has been proposed that A/T homopolymer tract expansion has a critical threshold of 7 to 10 bases in length, based on a study of the representation of homopolymer tracts in multiple organisms (17). Although somatic insertions do not necessarily reflect what occurs evolutionarily, it is not likely that the occurrence of strand slippage is different in germ cells and somatic cells. Our finding of over-representation of tract expansion at homopolymer lengths of 6 to 9 supports a narrow window of tract length for strand slippage occurrence. It is possible that the reannealing mispairing is more difficult due to the physical parameters, such as rigidity, of the DNA and less frequent DNA polymerase pausing for short homopolymer tracts.

About 1.55% (77 novel sequence insertions in a total of 4,972 unique insertions) of the naturally occurring unique insertion events are clearly not due to strand slippage during DNA synthesis because they bear no relation to the adjacent sequence on the same strand. These 77 events most likely occurred at sites of DNA double strand breaks (DSBs) with broken ends repaired by various enzymes. All except three events are insertions with ≤5-nt addition. Inspection of the inserted nucleotides, the surrounding sequences, and other DNA changes in proximity, shows that seven of the 77 DNA damage events are categorized as complex events. These events have additional somatic DNA changes, such as other insertion, deletion or base substitution, within a 10-nucleotide zone of the insertional event called by Mutect2 and Strelka (Supplementary Table S1). These are most likely NHEJ sites with complex processing of broken ends. The remaining 70 DNA damage events can be categorized into two groups (Supplementary Table S2). A key characteristic of Group 1 insertions is that all the consecutive nucleotides inserted can be found on the complementary strand adjacent to the insertion site. Group 1 insertions can be generated by a complementary strand nick and ligation mechanism (Fig. 7A and 7B). This proposed mechanism involves ligation of a DNA fragment released from a nick of the complementary DNA strand adjacent to the DSB site. Depending on whether endonuclease activity is present or whether non-templated nucleotide addition takes place at the DNA end, extra nucleotides may be present between the complementary sequence and the inserted nucleotides. Addition of nucleotides at DSB repair junctions have been observed in NHEJ reconstitution experiments containing only highly purified human NHEJ proteins and also in human extrachromosomal cellular assay systems (18–20). The observation of these general features of NHEJ in human colon crypts indicates that the observation in purified biochemical systems and mammalian cell lines resembles the naturally occurring NHEJ events, at least for this type of DNA breakage repair events. This categorization and the proposed mechanism can explain roughly half of the DNA breakage repair events involving insertion in a consistent and straightforward manner.

Group 2 DNA breakage repair events can only be explained by non-templated nucleotide addition by DNA polymerases, such as polμ and polα, to one of the broken DNA ends before joining of the two broken ends (21, 22). In Group 2, there are 31 insertions of 2- to 5-nt and three insertions of 13-, 15-, and 33-nt. The three long nucleotide additions would be unusual for polμ and polα template independent activity; more likely, the three long insertions may be the result of a random nuclear DNA fragment being ligated to the DSB site during repair.

Several possible mechanisms may explain the insertion of a single unit of duplicated sequence in a unique DNA region that leads to a new direct repeat. Similar to the nick and ligation process described above for short insertions, a DNA fragment released from a single strand nick adjacent to a DSB may be ligated to the same strand (not the complementary strand discussed above) of the other broken end. When new DNA synthesis fills in the gap, duplication of the released DNA fragment occurs (Fig. 7B). Alternatively, slippage during DNA synthesis can also lead to the same result. When a DNA polymerase pauses and the newly synthesized DNA strand becomes unpaired, microhomology at the 3’ end of the nascent strand and the template strand may mediate mispairing, permitting synthesis of a duplicated unit of DNA sequences (Fig. 7C). An insertion of this nature occurs in ∼ 30% of acute myeloid leukemia (AML) at the *FLT3* gene (23). The *FLT3* event may be triggered by nicking by an endogenous enzyme at the *FLT3* gene duplication sites.

The unique dataset we generated from 106 single human colon crypts with matched bulk DNA control provides the first opportunity to examine naturally occurring DNA breakage and the DNA strand slippage events in the human body without the limitations of cellular and biochemical experimental systems. We acknowledge that a perfectly restored DNA sequence at breakage sites cannot be identified in our dataset just as this limitation also applies to experiments in the cellular and biochemical systems. However, we are able to evaluate a relatively high number of DNA strand slippage and DNA breakage repair events and propose potential mechanisms to explain the DNA breakage repair events observed. The resemblance of the naturally occurring DNA breakage repair events to the cellular and biochemical systems provides useful confirmation across a range of detection methods. Events similar to the duplication in the *FLT3* gene allow for contemplation of potential mechanisms for disease causing mutations.

## ACKNOWLEDGEMENT

We would like to thank the Norris Comprehensive Cancer Center Translational Pathology Core for the sample collection. We would like to acknowledge the Norris Comprehensive Cancer Center Molecular Genomics Core and the Keck Genomics Platform at the University of Southern California for the sequencing work. This work is supported by funds from NIA R01 AG 067615 and the Catherine and Joseph Aresty Endowment to CLH. MRL was supported by NIGMS R35118009.

---

**Supplementary Figure S1.**
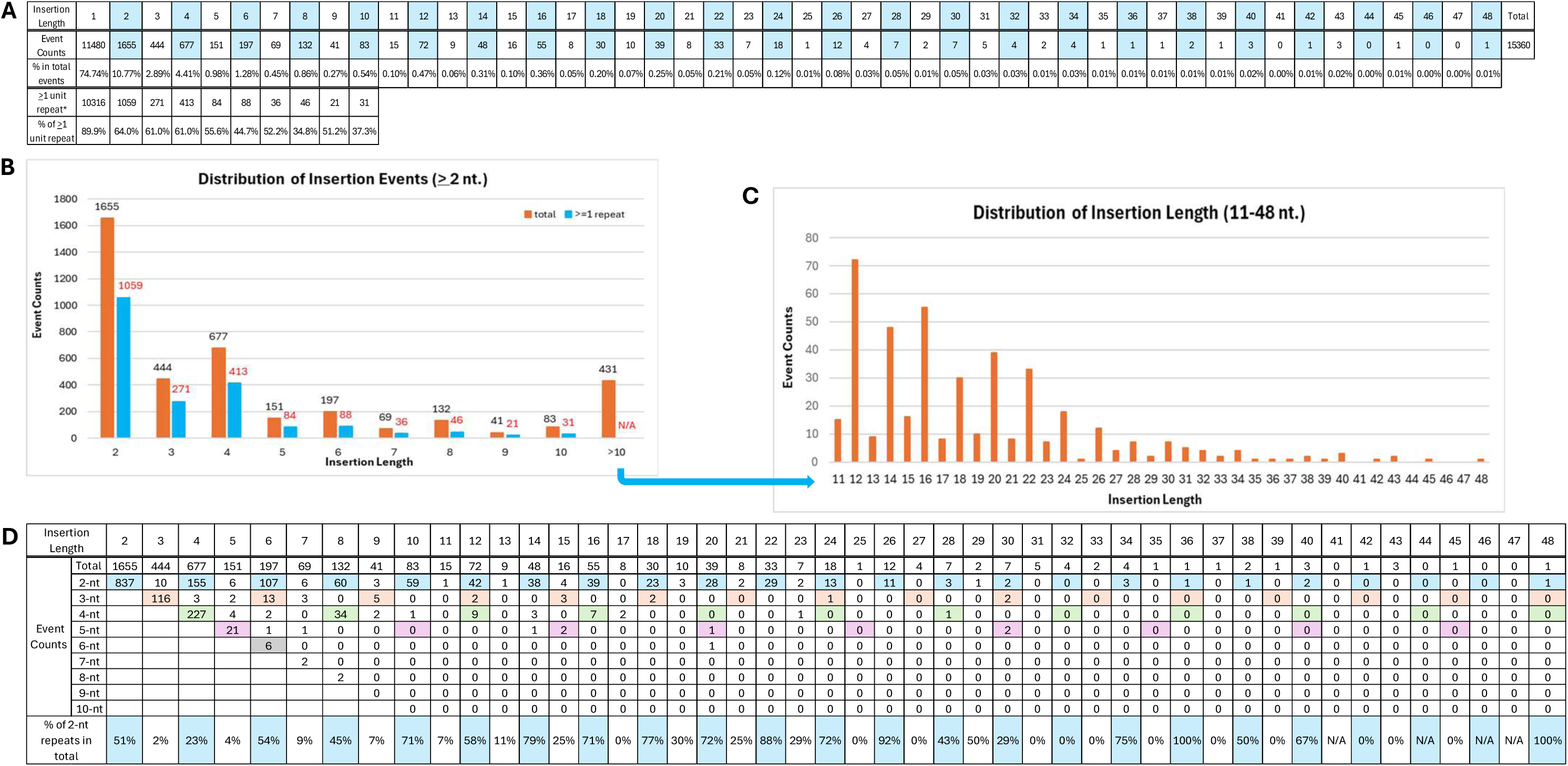
Distribution of Insertion Length from Preliminary Analysis of ALL dataset. A) Event count and the observed frequency in the total for each insertion length in the ALL dataset of 15,360 insertions from 106 single human colon crypts (21 individuals). The count of insertions with ≥1 unit of duplicated sequence in the adjacent region for insertion length 1 to 10 is also listed. Percentage of events with ≥1 unit of duplicated sequence in total occurrence of each insertion length is calculated. The blue highlight marks the even number insertion events. B) Graphical illustration of the event counts for insertion length from 2 to 10. C) Graphical illustration of the event counts for insertion length from 10 to 48. The event count is displayed on the top of each bar in B) and C). D) Simple sequence repeat (SSR, length of 2 to 10) with at least 2 units of repeat adjacent to each insertion length is identified and listed. The percentage of 2-nt SSR in the total for insertion length is calculated. The highlights are as indicated in Figure 1C.

**Supplementary Figure S2.**
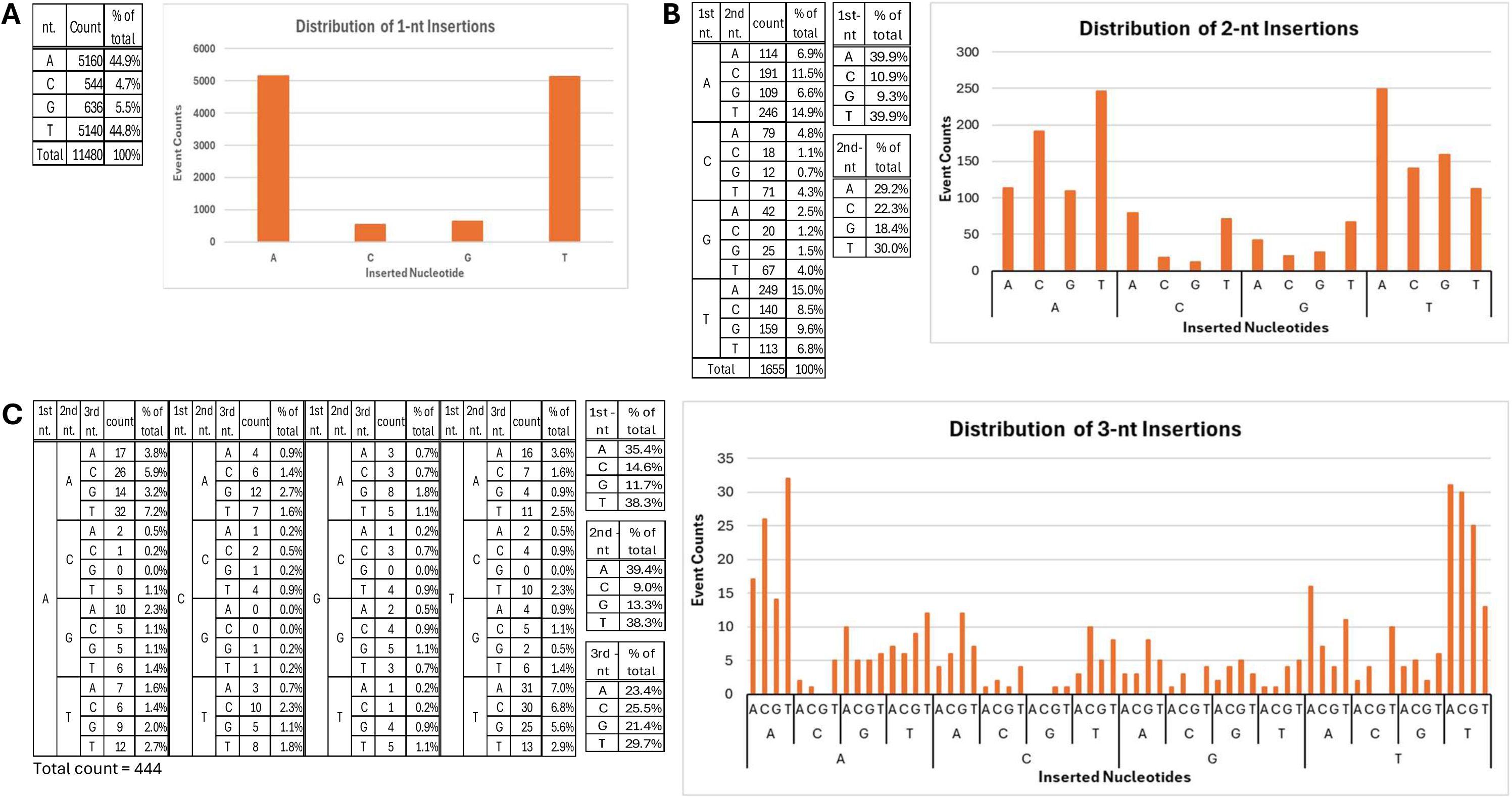
Base content analysis of one to three nucleotide insertions in the ALL dataset. A) Nucleotide content of single base (1-nt) insertions. B) Nucleotide content of two-base (2-nt) insertions. C) Nucleotide content of three-base (3-nt) insertions. The tabulation of nucleotides and percentage of each of the four nucleotides at each position of insertion is on the left and the graphical illustration of the nucleotide profile is on the right of each panel.

**Supplementary Figure S3.**
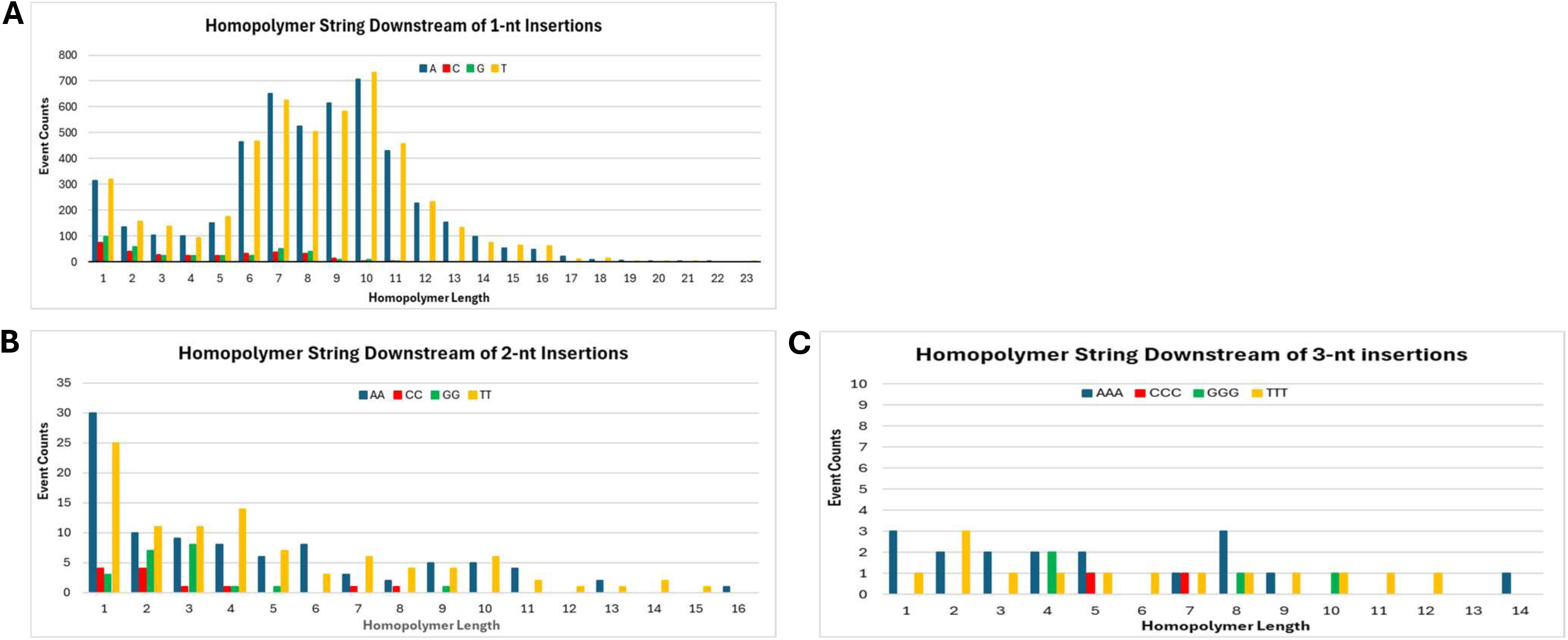
Presence of homopolymer at site of one- to three-nucleotide insertions in the ALL dataset. A) Event count of events with homopolymer length 1 to 23 at 1-nt insertion sites. Length of 0 indicates the 1-nt insertion is followed by a different nucleotide than the base inserted. B) Count of events with homopolymer length 0 to 9 at the site of 2-nt insertions with two identical nucleotides. C) Count of homopolymer length 0 to 6 at the site of 3-nt insertions with three identical nucleotides. The tabulation of events is above the graphic illustration in each panel. All lengths of homopolymer present in the dataset is included in the illustration.

## Notes

### Competing Interest Statement

The authors have declared no competing interest.

### Summary of Updates

correction of color marking of nucleotide in Figure 6B left panel

